# Mutations in the non-catalytic polyproline motif destabilize TREX1 and amplify cGAS-STING signaling

**DOI:** 10.1101/2024.01.04.574136

**Authors:** Abraham Shim, Xiaohan Luan, Wen Zhou, Yanick Crow, John Maciejowski

**Affiliations:** Molecular Biology Program, Sloan Kettering Institute, Memorial Sloan Kettering Cancer Center, New York, NY 10065, USA; Department of Immunology and Microbiology, School of Life Sciences, Southern University of Science and Technology, Shenzhen, Guangdong 518055, China; MRC Human Genetics Unit, Institute of Genetics and Cancer, The University of Edinburgh, Edinburgh, UK; Laboratory of Neurogenetics and Neuroinflammation, Imagine Institute, INSERM UMR1163, University Paris Cité, Paris, France

**Keywords:** cGAS, STING, TREX1, Aicardi-Goutières syndrome

## Abstract

The cGAS-STING pathway detects cytosolic DNA and activates a signaling cascade that results in a type I interferon (IFN) response. The endoplasmic reticulum (ER)-associated exonuclease TREX1 suppresses cGAS-STING by eliminating DNA from the cytosol. Mutations that compromise TREX1 function are linked to autoinflammatory disorders, including systemic lupus erythematosus (SLE) and Aicardi-Goutières syndrome (AGS). Despite key roles in regulating cGAS-STING and suppressing excessive inflammation, the impact of many disease-associated *TREX1* mutations - particularly those outside of the core catalytic domains - remains poorly understood. Here, we characterize a recessive AGS-linked TREX1 P61Q mutation occurring within the poorly characterized polyproline helix (PPII) motif. In keeping with its position outside of the catalytic core or ER targeting motifs, neither the P61Q mutation, nor aggregate proline-to-alanine PPII mutation, disrupt TREX1 exonuclease activity, subcellular localization, or cGAS-STING regulation in overexpression systems. Introducing targeted mutations into the endogenous *TREX1* locus revealed that PPII mutations destabilize the protein, resulting in impaired exonuclease activity and unrestrained cGAS-STING activation. Overall, these results demonstrate that TREX1 PPII mutations, including P61Q, impair proper immune regulation and lead to autoimmune disease through TREX1 destabilization.

## INTRODUCTION

Type I interferonopathies, such as the monogenic disease Aicardi-Goutières syndrome (AGS), often involve chronic systemic and neurological autoinflammation and high levels of type I interferon (IFN) activity in the blood and cerebrospinal fluid (Crow and Stetson, 2022). AGS can result from loss-of-function (or specific dominant-negative) mutations in *TREX1*, *RNASEH2A*, *RNASEH2B*, *RNASEH2C*, *SAMHD1*, and *ADAR1*, gain-of-function mutations in *IFIH1 (Lehtinen et al., 2008; Rice et al., 2007a)*. Mutations in *TREX1* are among the most common in AGS, accounting for nearly one-quarter of all AGS-linked mutations (Crow et al., 2015; Rice et al., 2007b).

TREX1 is a 3′→5′ exonuclease that degrades cytosolic DNA to act as a nucleolytic antagonist of the cGAS-STING pathway (Ablasser et al., 2014; Gray et al., 2015; Grieves et al., 2015; Mazur and Perrino, 2001; Stetson et al., 2008; Wolf et al., 2016). Binding to cytosolic DNA stimulates cGAS catalytic activity and the production of the 2′3′-cyclic GMP-AMP (cGAMP) second messenger (Ablasser et al., 2013; Diner et al., 2013; Gao et al., 2013). cGAMP engagement with its downstream receptor STING ultimately results in activation of the transcription factor IRF3 and the expression of type I IFNs and other immunomodulatory proteins (Ablasser and Chen, 2019).

Mouse models of TREX1 dysfunction recapitulate hallmarks of AGS and related disorders, including familial chilblain lupus. *Trex1*-deficient mice exhibit multi-organ inflammation and decreased survival (Grieves et al., 2015; Stetson et al., 2008). Replacement of the wild-type *Trex1* gene in mice with the nuclease-deficient *Trex1* D18N mutant results in a lupus-like disease (Grieves et al., 2015). The health and viability of *Trex1*-deficient animals are restored by deletion of *Cgas*, *Sting1*, *Irf3* and *Ifnar*, indicating that unchecked DNA sensing is responsible for the observed pathologies (Ablasser et al., 2014; Ahn et al., 2014; Gao et al., 2015; Gray et al., 2015; Stetson et al., 2008).

Specific dominant-negative mutations in *TREX1* include D18N, D200N, and H195Y, which disrupt key catalytic residues, and more frequently observed recessive alleles, e.g. R114H and R97H, that occur in the dimerization surface and hinder requisite homodimerization of TREX1 (Lehtinen et al., 2008; Rice et al., 2015). Other less-common mutations have been proposed to impede TREX1 function by altering phase separation or by destabilizing the protein (Zhou et al., 2022, 2021). The mechanisms associated with many disease-linked *TREX1* mutations are poorly understood.

Outside of its catalytic core, TREX1 possesses a single-pass transmembrane helix at its C-terminus that anchors the protein in the ER and positions the nuclease domain in the cytosol (Lee-Kirsch et al., 2007; Mazur and Perrino, 2001; Mohr et al., 2021; Wolf et al., 2016). Deleting this C-terminal extension ablates TREX1 ER localization but does not affect its catalytic activity (De Silva et al., 2007; Lee-Kirsch et al., 2007). *TREX1* mutations that truncate the C-terminus disrupt TREX1-ER association while preserving nucleolytic activity, and are associated with a distinct clinical disease referred to as retinal vasculopathy with cerebral leukoencephalopathy (RVCL) (Crow and Manel, 2015; Yan, 2017). RVCL is inherited in an autosomal dominant manner and lacks clear links to excessive type I IFN production (Rodero et al., 2017).

The non-repetitive proline-rich region termed the polyproline II helix (PPII) is another unique motif present in TREX1, but not found in other nucleases within the larger DnaQ family, including the closely related TREX2 homolog (Brucet et al., 2007; De Silva et al., 2007). Like the TREX1 C-terminal extension, the positioning of the PPII helix distal to the TREX1 active site and its absence from the otherwise closely related, catalytically active TREX2 nuclease suggest that it is also unlikely to participate in catalysis or DNA binding. The functional significance of this domain is not known.

Here, we report that TREX1 P61Q mutations located in the PPII motif are linked with AGS and show how these mutations destabilize TREX1 without directly affecting nucleolytic activity or subcellular localization. We demonstrate that TREX1 P61Q instability causes overactive cGAS-STING signaling, ultimately resulting in cGAMP overproduction and excessive levels of type I IFN expression. Thus, these results indicate that the TREX1 P61Q mutations cause AGS through TREX1 protein destabilization and suggest that protein destabilization may account for a subset of AGS patients with *TREX1* mutations.

## RESULTS

### TREX1 PPII mutations are associated with AGS

We identified proline-to-glutamine (P61Q) point mutations in *TREX1* in two patients from two families presenting with features of AGS (Fig. 1A; AGS972: c.182C>A p.Pro61Gln homozygote; AGS1583: p.Pro61Gln/Arg114His compound heterozygote) (Rice et al., 2017). Since R114H renders the allele null by disrupting obligate dimerization of TREX1 (Lehtinen et al., 2008), these findings suggest a recessive, loss-of-function nature of the P61Q mutation. Indeed, calculation of IFN scores, derived by measuring the expression of six IFN stimulated genes (ISGs) using quantitative polymerase chain reaction, revealed a significant upregulation of IFN signaling relative to persons considered to be controls, thus placing both individuals within the type I interferonopathy spectrum (Crow and Manel, 2015; Rice et al., 2017).

**Figure 1.**
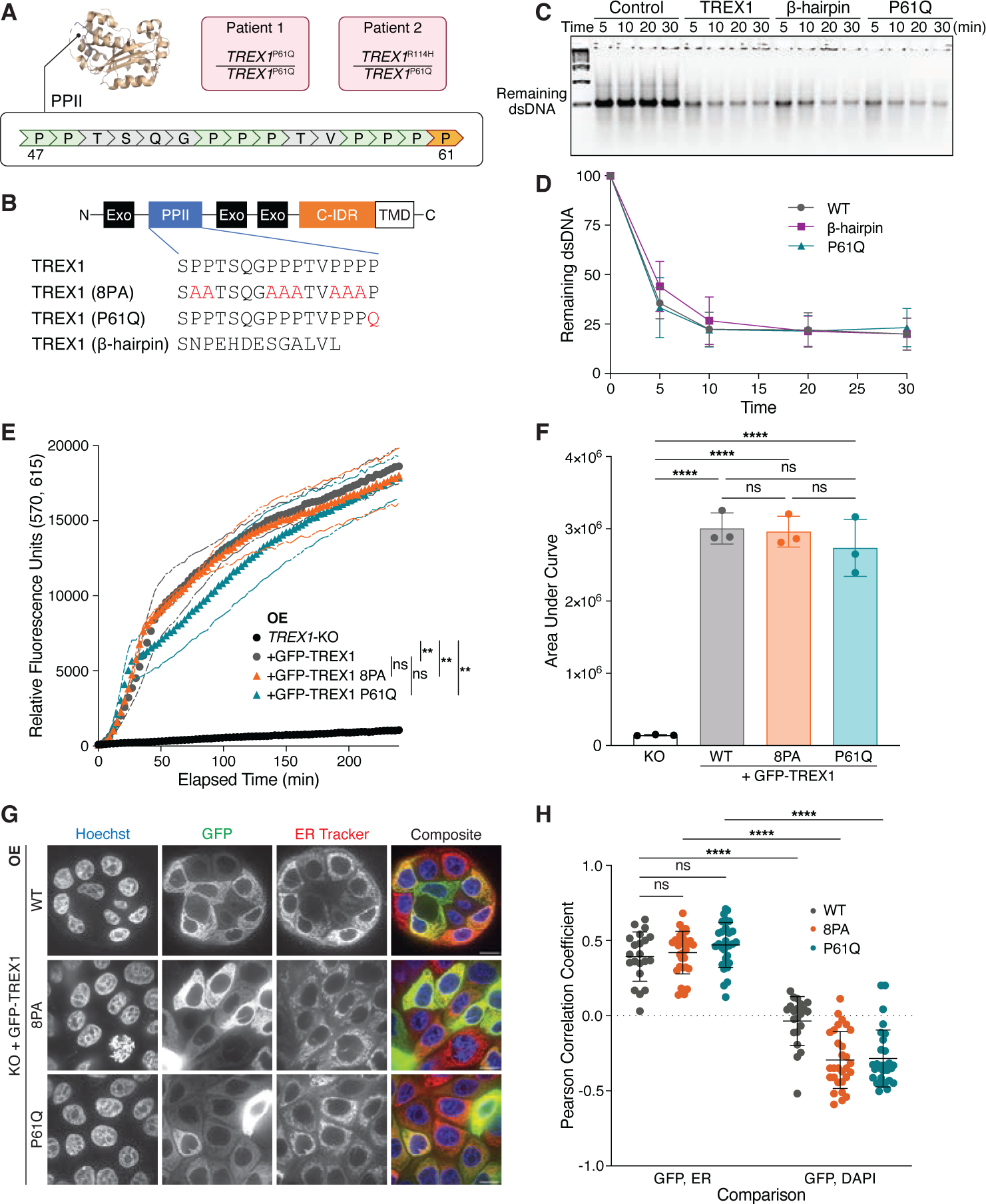
Mutations in PPII are linked to AGS but do not compromise intrinsic functions of overexpressed TREX1. **A.** Location of PPII and P61 (orange) within TREX1. Genotypes of two AGS patients harboring the P61Q mutation are shown in pink. **B.** Schematic of GFP-TREX1 mutants used to reconstitute MCF10A *TREX1* KO cells via lentiviral overexpression. Exo = exonuclease domain; C-IDR = C-terminal intrinsically disordered region; TMD = transmembrane domain responsible for TREX1-ER linkage. **C.** Representative DNA gel from *in vitro* nuclease assay. A dsDNA substrate was co-incubated with purified TREX1 mutant protein for the indicated duration. Control = no TREX1 added; β-hairpin = TREX1 with PPII replaced with TREX2 β-hairpin occurring at corresponding position as TREX1. P61Q = TREX1 P61Q **D.** Quantification of the *in vitro* nuclease assay in (C); mean ± s.d., *n* = 3, two-way ANOVA (interaction *p* = 0.5020). **E.** Time course fluorescence reading of the lysate-based nuclease assay. Briefly, a dsDNA substrate labeled with adjacent TEX615 fluorophore and Iowa Black quencher was co-incubated with whole cell lysates. 3′→5′ exonuclease activity eliminates the quencher, liberating TEX615 fluorescence; mean ± s.d., *n* = 3, \*\**p* < 0.01, ns = not significant, two-way ANOVA (interaction *p* < 0.0001, time *p* < 0.0001, genotype *p* < 0.0001). **F.** Definite integral values from *t* = 0 min to *t* = 240 min for each time course sample in (E); mean ± s.d., *n* = 3, \*\*\*\**p* < 0.0001, ns = not significant, one-way ANOVA (*p* < 0.0001). **G.** Live-cell images of GFP-TREX1 (green) in *TREX1*-KO cells. DNA was stained with Hoechst 33342 (blue) and ER was stained with ER Tracker Red (red). Scale bars = 10 μm. **H.** Pearson correlation coefficients of the indicated cells as in Fig. 1G; mean ± s.d., *n* = 5 experiments, \*\*\*\**p* < 0.0001, ns = not significant, two-way ANOVA (interactions *p* < 0.0001, comparison pair *p* < 0.0001, genotype *p* = 0.0027).

**Figure S1.**
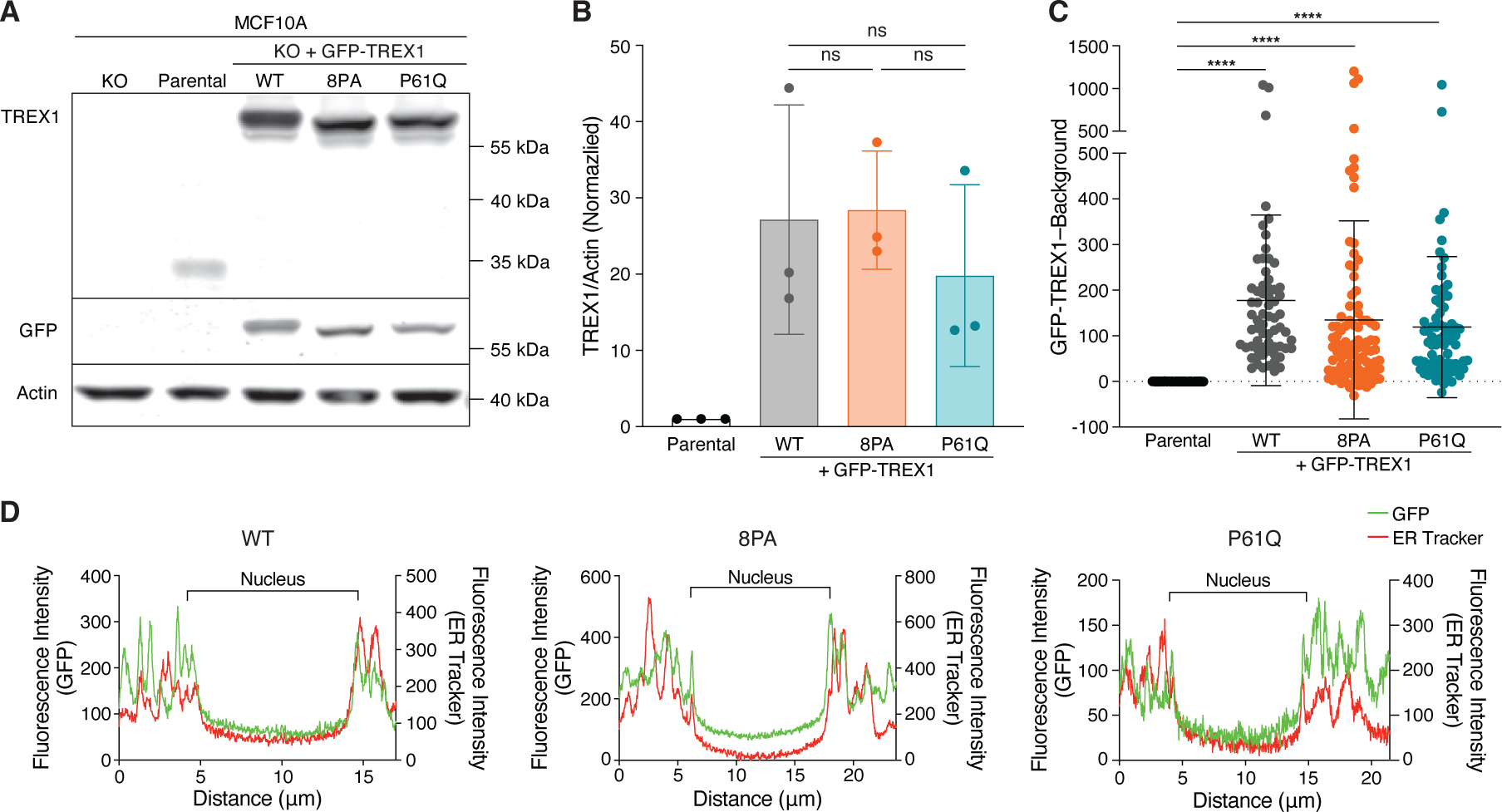
Overexpressed TREX1 mutants localize to the ER. **A.** Representative immunoblot of MCF10A *TREX1* KO cells reconstituted with the indicated GFP-TREX1, using anti-TREX1, anti-GFP, and anti-actin antibodies. **B.** Quantification of TREX1 immunoblot signal normalized to actin; mean ± s.d., *n* = 3, ns = not significant, one-way ANOVA (*p* = 0.0401). For each replicate, the parental TREX1/Actin signal was set to one. **C.** Quantification of GFP-TREX1 signal in the indicated cells as in Fig. 1G; mean ± s.d., *n* = 5 experiments, \*\*\*\**p* < 0.0001, one-way ANOVA (*p* < 0.0001).**D.** Line profile analysis of the indicated cells as in Fig. 1G. Position of the nucleus was determined using the line profile signal of the DAPI channel. Background signal was subtracted from all points.

Pro-61 lies in a proline-rich tract termed the polyproline II (PPII) helix (Fig. 1A,B) (Brucet et al., 2007; De Silva et al., 2007). PPII positioning distal to the TREX1 active site and its absence from other catalytically proficient enzymes of the DnaQ family, including TREX2, suggest it is likely to be dispensable for nucleolytic activity. To test this directly, we purified the N-terminal enzymatic domain of TREX1 proteins, including human TREX1, a TREX1 P61Q mutant, and a TREX1 PPII>β-hairpin chimera, in which the TREX1 PPII helix is replaced by the β-hairpin found in the corresponding position within TREX2 (Fig. 1B). As expected, *in vitro* nuclease assays using purified proteins demonstrated that TREX1 P61Q and β-hairpin mutants digested dsDNA with efficiencies comparable to the wild-type enzyme with >50% of substrate degraded within the first 5 minutes of incubation (Fig. 1C,D).

To further investigate the potential impact of TREX1 PPII mutations we assayed TREX1 exonuclease activity in cell lysates. In brief, lysates were incubated with a dsDNA substrate possessing a fluorescent label at one 5′ end closely positioned next to a 3′ quencher (Methods). TREX1 3′→5′ exonuclease activity is predicted to liberate the fluorescent dye from the 3′ quencher and thus result in the acquisition of fluorescence. Cell lysates were prepared from *TREX1*-deficient MCF10A cells stably transduced with GFP-TREX1-WT, GFP-TREX1-P61Q, and GFP-TREX1-8PA, in which eight prolines in PPII - excluding P61 - are mutated to alanine (Fig. 1B). Lentiviral transduction of these constructs into *TREX1*-deficient MCF10A cells yielded stable overexpression of GFP-tagged mutant proteins, with no significant differences in protein levels between the three genotypes (Fig. S1A and S1B). As expected, incubation of the dsDNA probe with lysates prepared from MCF10A *TREX1* KO cells reconstituted with GFP-TREX1-WT resulted in the rapid acquisition of fluorescence (Fig. 1E,F). In contrast, *TREX1* deletion severely diminished the acquisition of fluorescence, confirming the specificity of this assay for TREX1 exonuclease activity (Fig. 1E,F). Similar to results obtained using isolated proteins, measurement of GFP-TREX1-8PA and GFP-TREX1-P61Q activities exhibited no significant differences from GFP-TREX1-WT (Fig. 1E,F). Taken together, these data indicate that targeted mutations within the PPII helix do not directly interfere with TREX1 exonuclease activity and suggest that the PPII helix is dispensable for TREX1 exonuclease activity against dsDNA.

We previously demonstrated that TREX1 association with the ER is critical for processing a subset of cytosolic DNA substrates including nuclear aberrations like micronuclei (Mohr et al., 2021). Positioning of the PPII within the catalytic core and distal to the ER transmembrane domain at the C-terminus of TREX1 suggested that the PPII domain is likely dispensable for ER association. To test this possibility directly, we performed live-cell imaging of cells overexpressing GFP-TREX1 mutants to characterize their subcellular localization. As previously reported (Mohr et al., 2021; Stetson et al., 2008; Wolf et al., 2016), GFP-TREX1-WT was excluded from the nucleus and its localization significantly overlapped with the ER, as indicated by staining with an ER tracker dye (Fig. 1G,H). GFP-TREX1-8PA and GFP-TREX1-P61Q subcellular localizations could not be distinguished from that of the wild-type enzyme, suggesting that the PPII is dispensable for directing TREX1 ER association (Fig. 1G,H; Figure S1C,D). Together, these data indicate that PPII mutations are unlikely to cause TREX1 dysfunction by interfering with its ER localization.

### Overexpressed TREX1 PPII mutants suppress cGAS-STING signaling

To test whether PPII mutations affect cGAS activation, we quantified intracellular cGAMP via ELISA (Fig. 2A). MCF10A cells lack high levels of cytosolic DNA and do not show strong cGAS activity at baseline, even upon *TREX1* deletion (Mohr et al., 2021; Zhou et al., 2021). We therefore stimulated cGAS activation by herring testes (HT-) DNA transfection. ELISA analysis revealed low to undetectable amounts of cGAMP (0.3005±0.03630 s.d. fmol/µg protein) in MCF10A cells after HT-DNA stimulation (Fig. 2A). As expected, cGAMP levels increased dramatically in *TREX1* KO cell lysates following HT-DNA transfection (2.792±0.3207 s.d. fmol/µg protein) (Fig. 2A). Reconstitution of MCF10A *TREX1* KO cells by overexpressing GFP-TREX1-WT diminished cGAMP to levels observed in the parental cell line (0.4398±0.1119 s.d. fmol/µg protein) (Fig. 2A). In keeping with their catalytic proficiency and normal ER localization, GFP-TREX1-8PA and GFP-TREX1-P61Q overexpression led to cGAMP reductions that were comparable to the wild-type GFP-TREX1 transgene (0.7678±0.2449 s.d. fmol/µg protein for GFP-TREX1-8PA; 0.9510±0.06908 s.d. fmol/µg protein for GFP-TREX1-P61Q).

**Figure 2.**
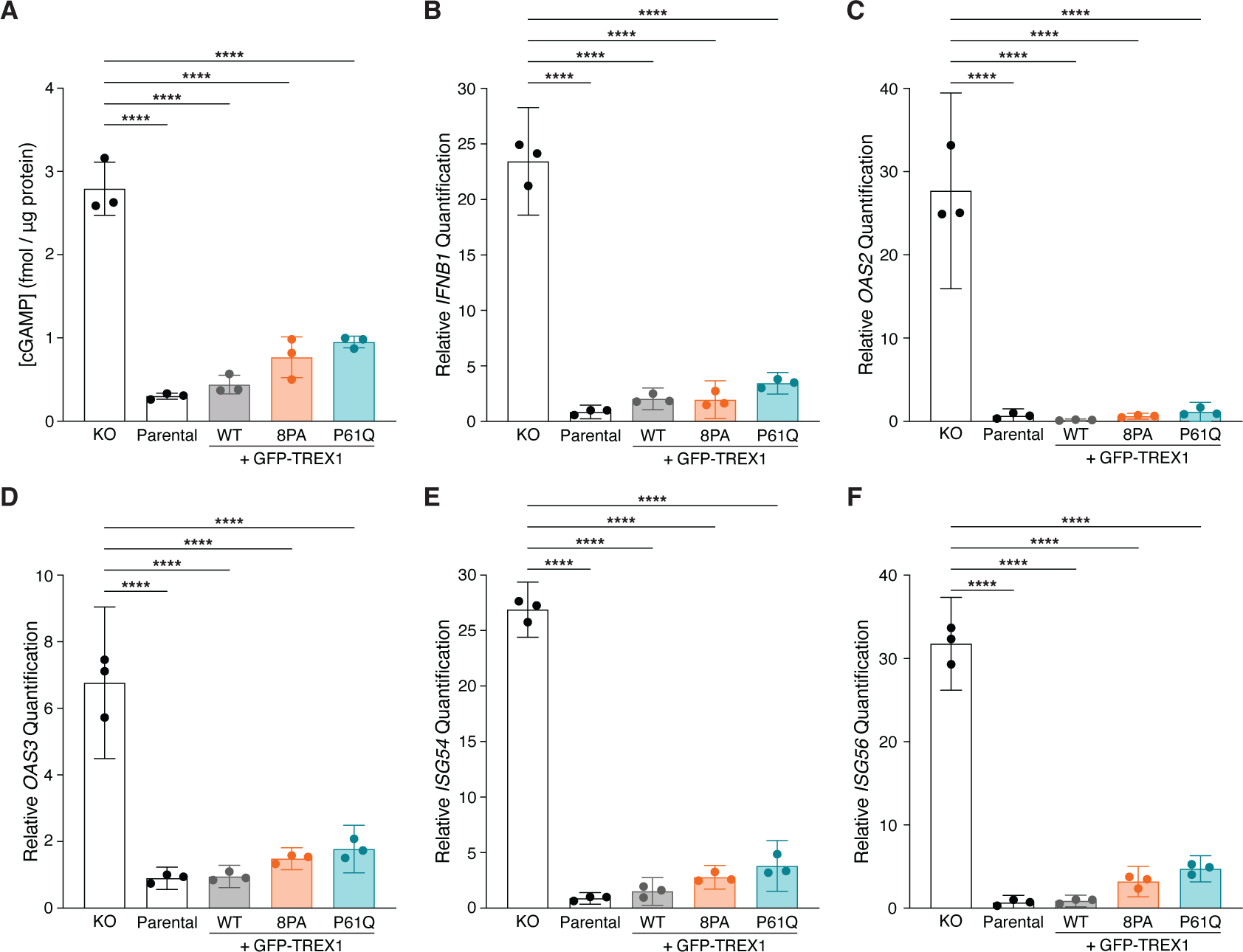
Overexpressed TREX1 mutants can suppress cGAS-STING signaling. **A.** ELISA analysis of cGAMP production in the indicated cells following the transfection of 4 μg HT-DNA; mean ± s.d., *n* = 3, \*\*\*\**p* < 0.0001, one-way ANOVA (*p* < 0.0001). **B–F.** RT-qPCR of *IFNB1*, *OAS2*, *OAS3*, *ISG54*, and *ISG56* expression in the indicated MCF10A cells following the transfection of 4 μg HT-DNA; mean ± s.d., *n* = 3, \*\*\*\**p* < 0.0001, one-way ANOVA (*p* < 0.0001 for *IFNB1*, *p* < 0.0001 for *OAS2*, *p* < 0.0001 for *OAS3*, *p* < 0.0001 for *ISG54*, and *p* < 0.0001 for *ISG56*).

**Figure S2.**
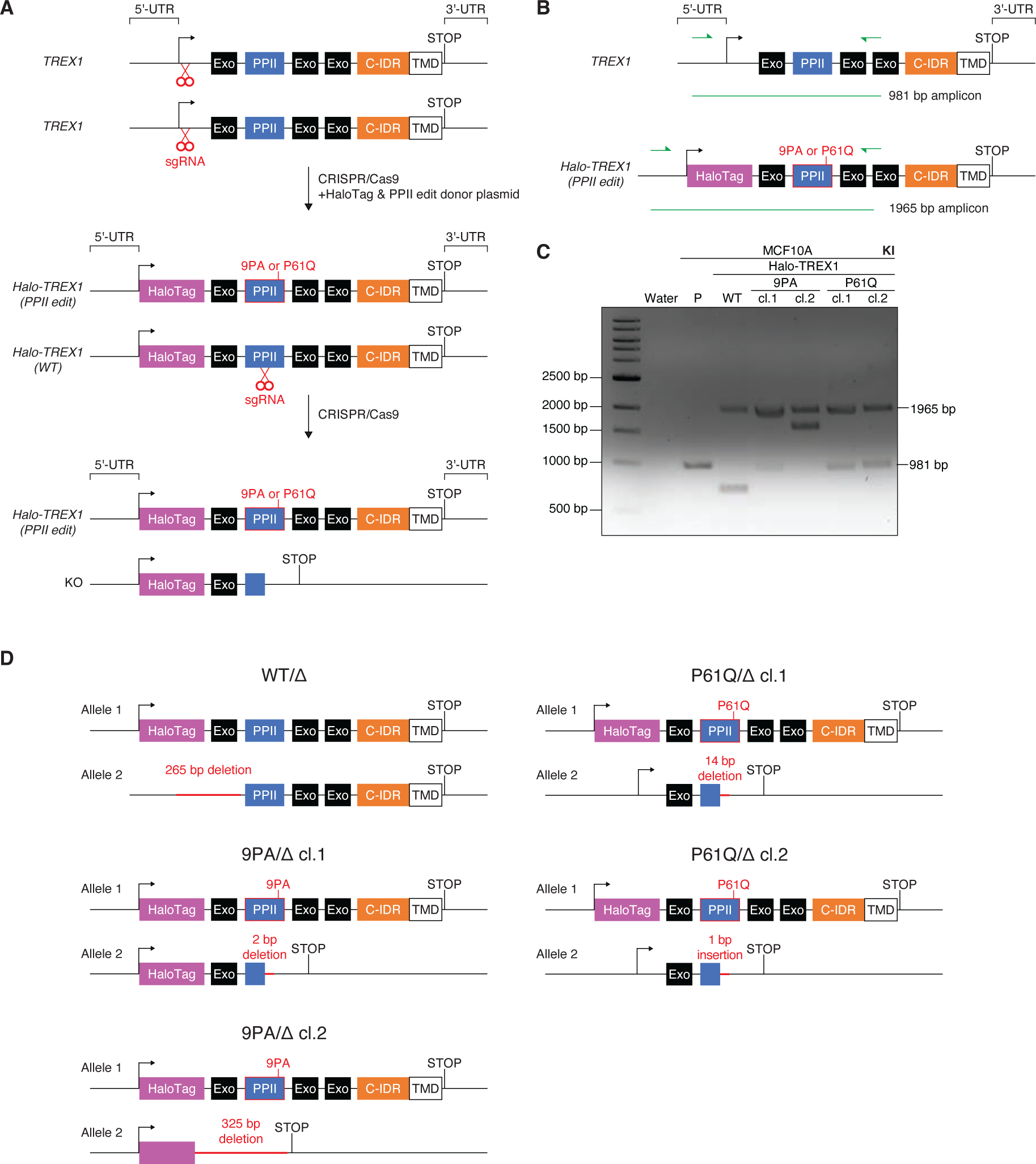
Generation of *TREX1* knock-in mutations. **A.** Representative schematic of *TREX1* gene editing protocol. Briefly, an N-terminal sgRNA and a HaloTag donor plasmid harboring a PPII edit in its downstream homology arm were nucleofected into MCF10A cells. Sanger sequencing revealed that HaloTag insertion occurs more frequently than incorporation of the PPII edit, often yielding two Halo-tagged alleles, one with the desired PPII edit. A second round of gene editing was carried out using a sgRNA specific for unedited PPII, knocking out the unedited allele while leaving the PPII-edited, Halo-tagged allele intact. **B.** Schematic of PCR primers and amplicons used to validate knock-in cell lines. **C.** PCR gel of all knock-in cell line clones used in this manuscript. All bands were excised and Sanger sequenced to validate expected gene edits. **D.** Schematic detailing the precise edits present in all clones.

We next sought to determine if TREX1 PPII mutations impacted the downstream cGAS-STING response by using RT-qPCR to measure expression of *IFNB1* and interferon-stimulated genes (ISGs) such as *OAS2*, *OAS3*, *ISG54*, and *ISG56* (Fig. 2B-F). As expected, RT-qPCR revealed strong increases in *IFNB1* and ISG mRNA levels in *TREX1* KO MCF10A cells upon HT-DNA stimulation relative to parental controls (Fig. 2B-F). In line with our cGAMP ELISA results, GFP-TREX1-WT, GFP-TREX1-8PA and GFP-TREX1-P61Q suppressed *IFNB1* and ISG expression to similar degrees upon overexpression in *TREX1* KO cells (Fig. 2B-F). Thus, counter to expectations based on the association between the TREX1 P61Q mutations and AGS (Fig. 1A), these results indicate that TREX1 PPII mutants are functionally proficient to suppress cGAS activation and downstream ISG expression upon overexpression in MCF10A cells.

### TREX1 PPII mutations destabilize the protein

We reasoned that strong TREX1 overexpression resulting from lentiviral delivery (Fig. S1A and S1B) may obscure defects associated with PPII mutation. We therefore used CRISPR-Cas9 gene editing to endogenously introduce an N-terminal HaloTag concurrently with a PPII edit—proline-to-alanine mutation of all nine prolines in PPII (9PA) or P61Q - into the diploid MCF10A cell line (Fig. S2A). The remaining, unedited allele was deleted, yielding Halo-TREX1/Δ genotypes for all subsequent experiments (Fig. S2A). All gene edits were validated by Sanger sequencing and PCR screening (Fig. S2B–D). Immunoblotting with anti-TREX1 antibodies further confirmed successful insertion of the HaloTag into the endogenous *TREX1* locus. (Fig. 3A).

**Figure 3.**
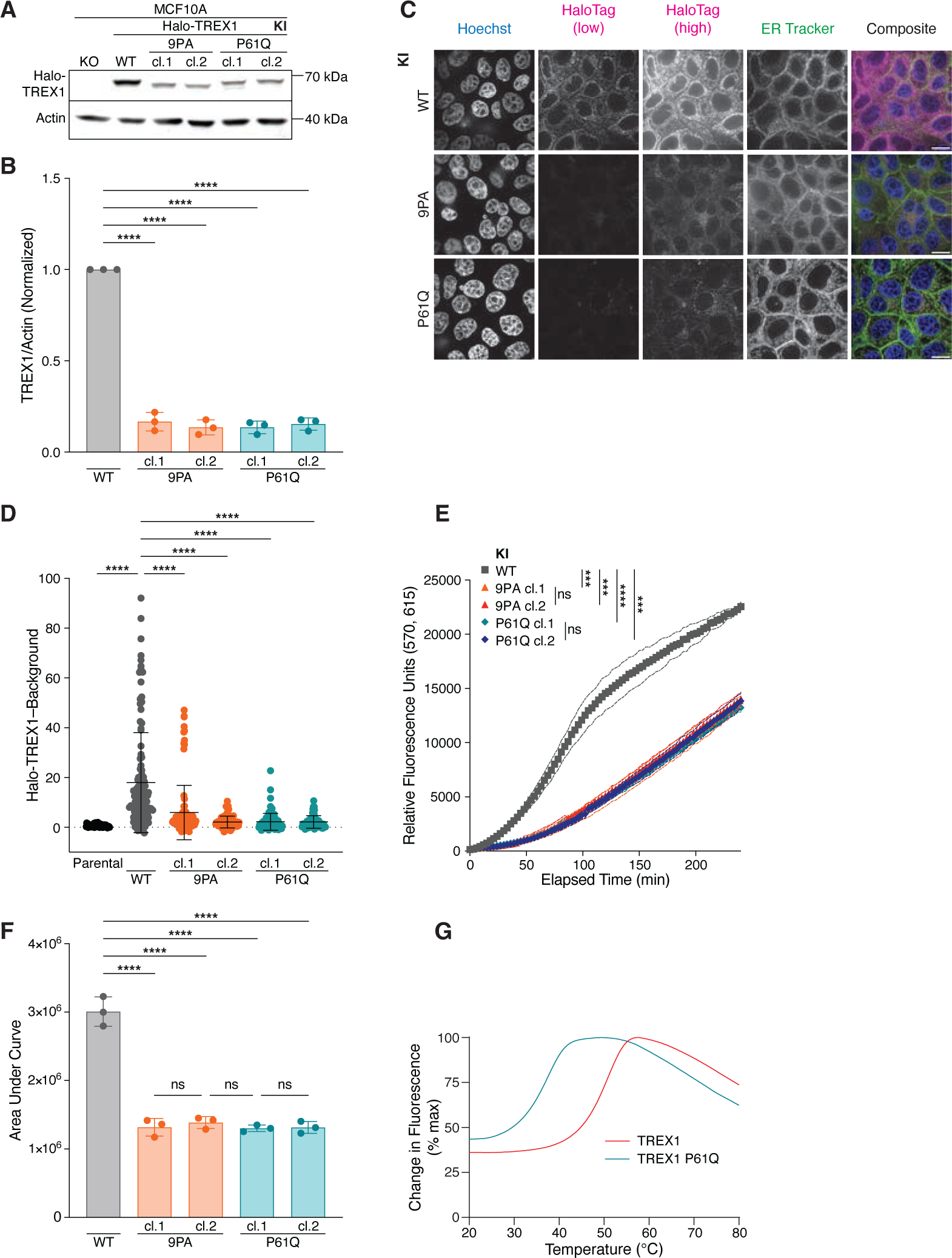
PPII mutations destabilize TREX1 and reduce TREX1 exonucleolytic activity. **A.** Immunoblot of MCF10A knock-in (KI) cell lines using anti-TREX1 and anti-actin antibodies. **B.** Quantification of TREX1 immunoblot signal normalized to actin; mean ± s.d., *n* = 3, \*\*\*\**p* < 0.0001, one-way ANOVA (*p* < 0.0001). For each replicate, the WT TREX1/Actin signal was set to one. **C.** Live-cell images Halo-TREX1 (magenta) in MCF10A knock-in cell lines. DNA was stained with Hoechst 33342 (blue) and ER was stained with ER Tracker Green (green). Halo-TREX1 images are shown using two lookup tables in order to highlight differences in fluorescence signal (HaloTag low) and to depict ER localization (HaloTag high). Scale bars = 10 μm. **D.** Quantification of Halo-TREX1 signal in the indicated MCF10A cells as in (E); mean ± s.d., *n* = 5 experiments, \*\*\*\**p* < 0.0001, one-way ANOVA (*p* < 0.0001). **E.** Time course fluorescence reading of lysate-based nuclease assay, using MCF10A knock-in cell lines; mean ± s.d., *n* = 3, \*\*\**p* < 0.001, \*\*\*\**p* < 0.0001, ns = not significant, two-way ANOVA (interaction *p* < 0.0001, time *p* < 0.0001, genotype *p* < 0.0001). **F.** Definite integral values from *t* = 0 min to *t* = 240 min for each time course sample in (E); mean ± s.d., *n* = 3, \*\*\*\**p* < 0.0001, one-way ANOVA (*p* < 0.0001). **G.** Thermal shift assay using purified TREX1 proteins. T_m_ = 51 °C for TREX1 WT, T_m_ = 37.5 °C for TREX1 P61Q.

**Figure S3.**
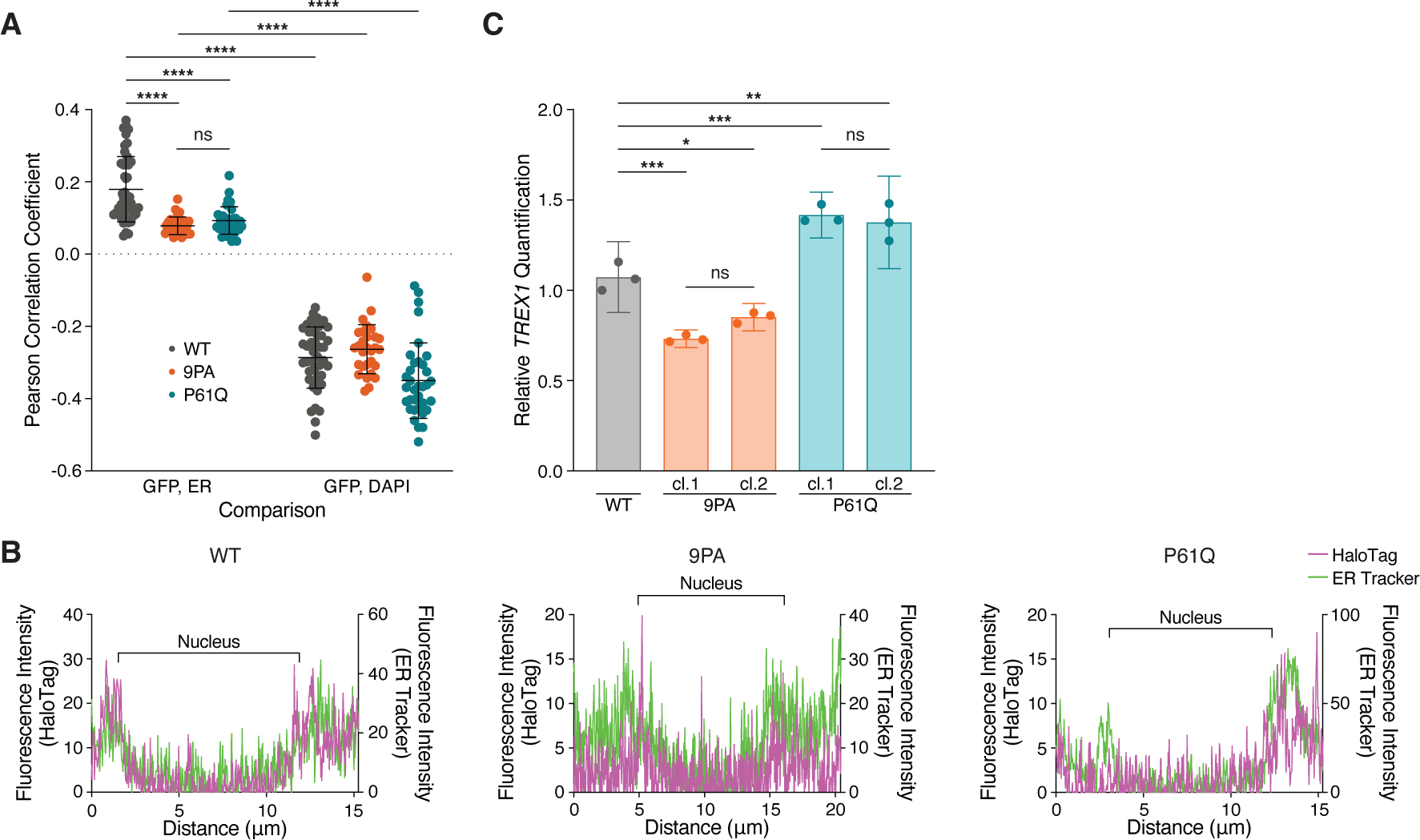
Mutations in PPII do not interfere with TREX1 transcription or localization. **A.** Pearson correlation coefficients of the indicated cells as in Fig. 3C; mean ± s.d., *n* = 5 experiments, \*\*\*\**p* < 0.0001, ns = not significant, two-way ANOVA (interaction *p* < 0.0001, comparison pair *p* < 0.0001, genotype *p* < 0.0001). **B.** Line profile analysis as indicated in Fig. 3C. Extent of the nucleus was determined using the line profile signal of the DAPI channel. Background signal was subtracted from all points. **C.** RT-qPCR of *TREX1* in the indicated cells following mock transfection; mean ± s.d., *n* = 3, \**p* < 0.05, \*\**p* < 0.01, \*\*\**p* < 0.001, ns = not significant, one-way ANOVA (*p* < 0.0001).

Interestingly, immunoblotting revealed significantly diminished Halo-TREX1-9PA and Halo-TREX1-P61Q signals in multiple, independently isolated subclones relative to the wild-type Halo-TREX1 control (Fig. 3A,B). Live-cell imaging confirmed decreased expression of Halo-TREX1-9PA and Halo-TREX1-P61Q relative to wild-type Halo-TREX1 (Fig. 3C and 3D). As expected, neither mutation compromised the ER localization of TREX1 (Fig. S3A and S3B). Similar to our prior results from GFP-TREX1 overexpression (Fig. 1E and 1F), all Halo-TREX1 lysates retained the ability to digest dsDNA (Fig. 3E and 3F). However, fluorescence increased at a much slower rate in Halo-TREX1-9PA/Δ and Halo-TREX1-P61Q/Δ lysates than Halo-TREX1-wild-type/Δ lysates, with the area under curve values decreased about two-fold. Thus, TREX1-P61Q and TREX1-9PA mutations lead to significant reductions in protein levels that are associated with corresponding decreases in nucleolytic activity.

Observed reductions in TREX1-9PA and TREX1-P61Q protein levels and activity could not be explained by reduced *TREX1* mRNA expression (Fig. S3C). Instead, Thermofluor analysis of purified TREX1 and TREX1-P61Q proteins demonstrated a significant 13.5°C difference in protein stability with TREX1 exhibiting a melting temperature (T_m_) of 51°C and TREX1-P61Q exhibiting a T_m_ of 37.5°C (Fig. 3G). These results indicate that TREX1 PPII mutations destabilize the protein, and thus lead to reduced overall protein levels with corresponding decreases in nucleolytic activity.

### cGAS-STING signaling is elevated in TREX1 PPII mutant cells

We next asked whether Halo-TREX1-PPII mutations interfere with cGAS-STING regulation. cGAMP ELISA analysis following stimulation by HT-DNA transfection demonstrated significant increases in intracellular cGAMP levels in multiple, independently isolated Halo-TREX1-9PA (1.092±0.1379 s.d. fmol/µg protein for clone 1; 1.161±0.06758 s.d. fmol/µg protein for clone 2) and Halo-TREX1-P61Q (1.171±0.1046 s.d. fmol/µg protein for clone 1; 0.8806±0.1024 s.d. fmol/µg protein for clone 2) mutant cell lines relative to a wild-type Halo-TREX1 control line (0.3194±0.07516 s.d. fmol/µg protein) (Fig. 4A). Indeed, cGAMP levels in lysates prepared from Halo-TREX1-P61Q and Halo-TREX1-9PA mutant cells more closely resembled levels measured in *TREX1* KO lysates (1.374±0.04191 s.d. fmol/µg protein). cGAMP levels were unchanged in all cell lines tested, including *TREX1* KO lines, following mock transfection, further confirming that MCF10A cells lack sufficient cytosolic dsDNA to activate an immune response under baseline conditions (data not shown).

**Figure 4.**
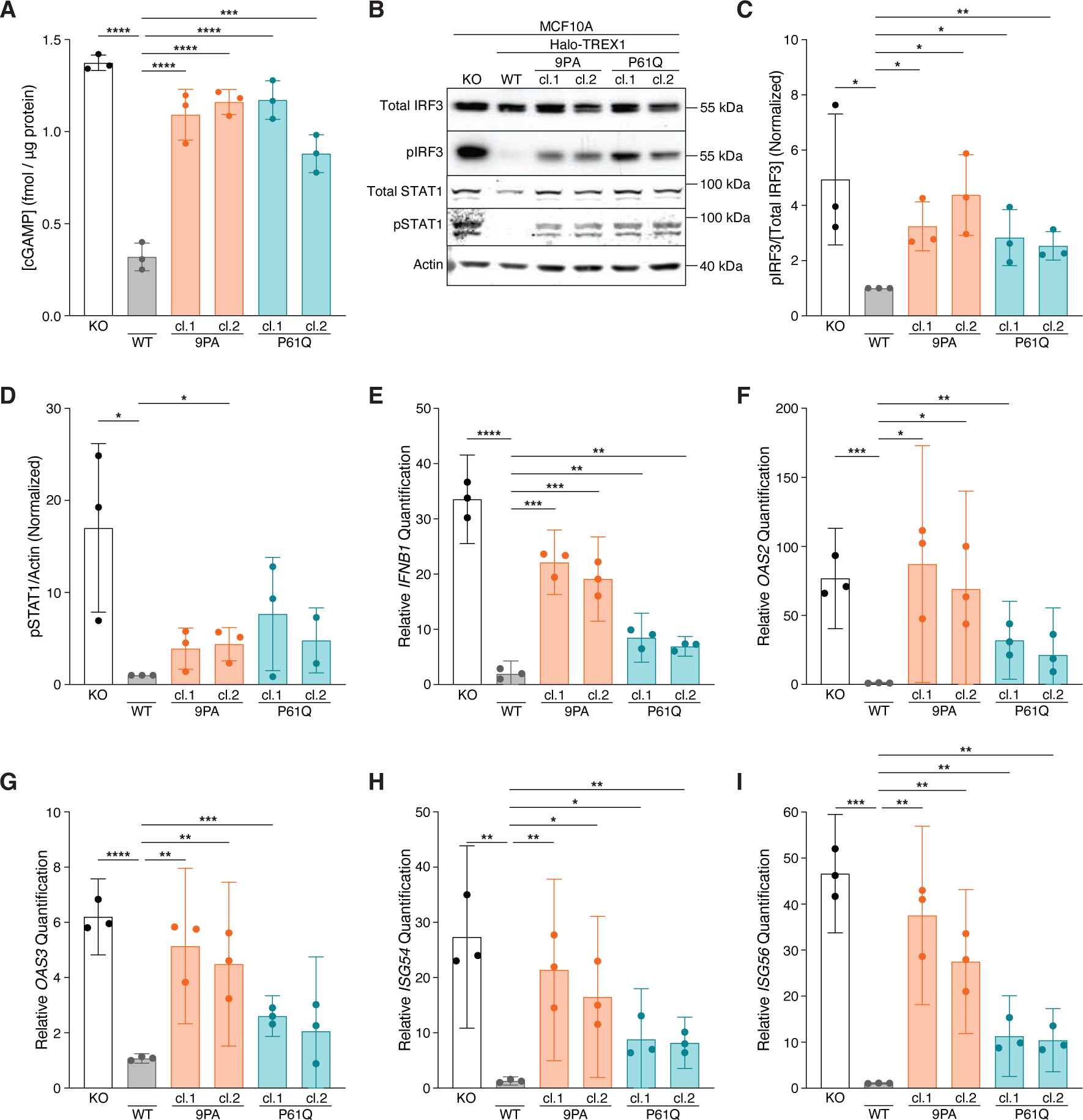
Mutations in PPII activate cGAS-STING signaling. **A.** ELISA analysis of cGAMP production in the indicated cells following the transfection of 4 μg HT-DNA; mean ± s.d., *n* = 3, \*\*\**p* < 0.001, \*\*\*\**p* < 0.0001, one-way ANOVA with post-hoc pairwise comparisons (*p* < 0.0001). **B.** Immunoblot of MCF10A knock-in (KI) cell lines using anti-IRF3, anti-phospho-S386 IRF3, anti-STAT1, anti-phospho-Y701 STAT1, and anti-actin. **C.** Quantification of phospho-S386 IRF3 immunoblot signal normalized to total IRF3 as in Fig. 4B; mean ± s.d., *n* = 3, \**p* < 0.05, \*\**p* < 0.01, unpaired two-tailed *t*-tests. For each replicate, the WT pIRF3/[Total IRF3] signal was set to one. **D.** Quantification of phospho-Y701 STAT1 immunoblot signal normalized to actin; mean ± s.d., *n* = 3, \**p* < 0.05, unpaired two-tailed *t*-tests. For each replicate, the WT pSTAT1/actin signal was set to one. **E–I.** RT-qPCR of *IFNB1*, *OAS2*, *OAS3*, *ISG54*, and *ISG56* expression in the indicated cells following the transfection of 4 μg HT-DNA; mean ± s.d., *n* = 3, \**p* < 0.05, \*\**p* < 0.01, \*\*\**p* < 0.001, \*\*\*\**p* < 0.0001, unpaired two-tailed *t*-tests.

Following cGAS-STING activation, TBK1 phosphorylates the transcription factor IRF3 at multiple residues including S386 and S396, inducing IRF3 dimerization and transcription of type I IFN (Liu et al., 2015). Increased type I IFN signaling results in the phosphorylation and activation of STAT1 (pY701)/STAT2 heterodimers, ultimately culminating in the transactivation of a wide-ranging pro-inflammatory response (Galluzzi et al., 2018). We therefore immunoblotted for phospho-IRF3 (pS386) and phospho-STAT1 (pY701) to assess cGAS-STING signaling downstream of cGAMP production (Fig. 4B-D). Consistent with prior work (Mohr et al., 2021), *TREX1* KO cells exhibited significant increases in the phosphorylated forms of IRF3 and STAT1 following HT-DNA stimulation relative to wild-type Halo-TREX1 controls (Fig. 4B-D). Congruent with the observed increase in cGAMP levels, Halo-TREX1-9PA and Halo-TREX1-P61Q mutant cells exhibited increased levels of pIRF3 and pSTAT1 compared to wild-type controls, albeit to a lesser extent than *TREX1* KO cells (Fig. 4B-D).

We next performed RT-qPCR to measure *IFNB1* and associated ISG mRNA levels to test if increases in cGAMP, and IRF3, and STAT1 phosphorylation are associated with elevated pro-inflammatory gene expression. Indeed, *IFNB1*, *OAS2*, *OAS3*, *ISG54*, and *ISG56* transcripts were elevated across multiple Halo-TREX1-9PA and Halo-TREX1-P61Q mutant subclones relative to wild-type controls to levels that were often indistinguishable from *TREX1* KO cells (Fig. 4E-I). Taken together, these results indicate that TREX1 PPII mutations result in defective cGAS regulation and an increased pro-inflammatory transcriptional response, defects most likely stemming from TREX1 protein instability and associated reductions in overall TREX1 protein levels and corresponding decreases in nucleolytic activity.

## DISCUSSION

Genetic associations of type I interferonopathies like AGS have been well-characterized, particularly in cases involving *TREX1* mutations linked with compromised catalytic activity (Crow and Manel, 2015). Yet, how missense mutations outside of the catalytic site can lead to inflammatory disease has often remained unclear. Here, we identify an AGS-linked P61Q point mutation within the non-catalytic PPII motif of TREX1. Using *in vitro* biochemical measures of protein stability and endogenous gene editing, we show that TREX1 PPII mutations, including P61Q, destabilize the protein, resulting in significantly decreased TREX1 protein levels, diminished TREX1 exonucleolytic activity, and impaired cGAS-STING regulation. These defects were obscured in lentiviral delivery models where massive overexpression of TREX1 PPII mutants masks reductions in protein stability to maintain effective cGAS inhibition. The distal position of PPII to the catalytic site, along with the lack of differences in the GFP-TREX1 lysate-based nuclease assay, suggests that the nucleolytic defect observed in the endogenous system is due to decreased protein levels, rather than a direct effect of the mutations on catalysis. Thus, our results indicate diminished protein stability and an associated reduction in overall nucleolytic power of TREX1 as a plausible molecular explanation for why TREX1 P61Q mutations lead to severe AGS phenotypes in patients.

Autoinflammatory disease-linked *TREX1* missense mutations often affect residues that play direct roles in DNA binding (i.e. R128H, K160R), catalytic activity (D18N/H, H195Y/Q, D200H/N) or dimerization (R97H, R114H). We recently reported that *TREX1* mutations may also cause dysfunction by interfering with TREX1 interactions with cGAS-DNA condensates (E198K) (Zhou et al., 2021). Here, the identification of AGS-linked TREX1 P61Q mutations suggests that another class of mutations may compromise TREX1 function by diminishing overall protein stability. Indeed, structural analyses predict that the disease-linked TREX1 T13N, T32R, R185C, and D220G substitutions are likely to diminish protein stability (Zhou et al., 2022). Biochemical experimentation supports this premise as TREX1 T13N, T32R, R185C, and D220G substituted proteins exhibit T_m_ reductions of 4–8 °C *in vitro* (Zhou et al., 2022). Thus, TREX1 protein destabilization may be a common defect occurring across multiple AGS-linked *TREX1* mutations.

TREX1 P61 is located with the PPII polyproline helix, a proline-rich region containing 9 prolines within a 15 amino acid stretch (Brucet et al., 2007; De Silva et al., 2007). This type of proline-rich segment is a conserved feature of TREX1, as it occurs in all organisms harboring TREX1, including placental mammals and marsupials. The paralog TREX2, as well as the ancient TREX nuclease occurring in non-mammals such as *Anopheles* and *Drosophila*, lack a proline-rich motif (Brucet et al., 2007), indicating that PPII likely evolved during the gene duplication event. Interestingly, the emergence of PPII in evolution seems to have coincided with the addition of a long C-terminal intrinsically disordered region. In keeping with structure-based predictions based on the PPII positioning outside of the catalytic core and ER transmembrane domains, our data confirm that the PPII motif is dispensable for TREX1 nucleolytic activity and subcellular localization. The precise function of the PPII motif therefore remains unknown.

The close positioning of the two PPII helices along the same side of the TREX1 dimer interface has been proposed to create a surface that allows for protein-protein interactions without occluding the active sites (Brucet et al., 2007; De Silva et al., 2007). Indeed, their high potential for presenting exposed hydrogen bond donors and acceptors, cause proline-rich motifs to be considered likely protein interaction domains (Adzhubei et al., 2013). The amino acid sequence of PPII matches the binding motif for the WW domain (Brucet et al., 2007), a peptide module characterized by two tryptophan residues (Sudol et al., 1995). Co-immunoprecipitation experiments have previously confirmed that murine TREX1 PPII interacts with the WW domain protein CA150 *in vitro* (Brucet et al., 2007). Whether human TREX1 also interacts with WW domain proteins and endogenous interactors remains unknown. Outside of a proposed interaction with the nucleosome assembly SET protein (Chowdhury et al., 2006), TREX1 protein partners are largely uncharacterized. Further work is therefore necessary to investigate this exciting hypothesis.

Our study relies heavily on the N-terminal HaloTag for studying the behavior of endogenous *TREX1* PPII mutations. We observed an apparent stabilizing effect of the HaloTag on TREX1, as Halo-TREX1(WT)/Δ yielded a stronger immunoblot signal than parental cells (data not shown). This observation is consistent with a prior report, which demonstrated that HaloTags can elicit a significant impact on the detection of proteins by Western blot (Broadbent et al., 2023). Apparent increases of HaloTag protein levels were attributed to enhanced western blot transfer efficiency (Broadbent et al., 2023). Therefore, western blotting analysis may underestimate the full extent of TREX1 P61Q protein instability. A further potential limitation of our study is the use of the non-malignant MCF10A breast epithelial cell line to model AGS-linked *TREX1* mutations. MCF10A cells were selected for this study because they possess an intact cGAS-STING-TREX1 pathway (Mohr et al., 2021) and are suitable for facile gene editing. However, it is not clear how well this cell model recapitulates aspects of AGS, a disease that primarily affects the central nervous system. Nevertheless, orthogonal measurements of TREX1 P61Q stability via Thermofluor analysis provide assurance that the P61Q mutation is likely to exert a destabilizing effect across multiple cell types and thus reinforce our proposed mechanism of pathogenesis in patients harboring the TREX1 P61Q mutation.

## METHODS

### Experimental Model and Subject Details

MCF10A cells were cultured in a 1:1 mixture of F12:DMEM media, supplemented with 5 % horse serum (Thermo Fisher Scientific #26050088), 20 ng/mL human EGF (Sigma Aldrich #E9644-.2mg), 0.5 mg/mL hydrocortisone (Sigma Aldrich), 100 ng/mL cholera toxin (Sigma Aldrich #H0888), 10 μg/mL recombinant human insulin (Sigma Aldrich #I9278-5ml), and 1% penicillin-streptomycin (Thermo Scientific #15140122). All media were supplied by the MSKCC Media Preparation core facility.

For HaloTag insertion and PPII gene editing of endogenous *TREX1* in MCF10A cells, an RNP mix was prepared by mixing 10 μg purified SpCas9 and 500 pmol of sgRNA (TREX1_gRNA#1, see Key Resources Table). After a 10-minute incubation at room temperature, 2500 ng of the pUC19-HA-Halo-TREX1 plasmid harboring the desired mutation was added to the RNP mix. The RNP-plasmid mixture was nucleofected using 4D-Nucleofector X Unit (Lonza). Fluorescence-activated cell sorting was used to isolate single-cell clones from the polyclonal cell population. For monoallelic knockout of *TREX1*, a pUC19-BBsI-CBh-TREX1_gRNA#2-Cas9-T2A-mCherry plasmid (Mohr et al., 2021) was transfected using Lipofectamine 3000 (Invitrogen #L3000075). Single-cell clones were isolated by limiting dilution culture.

### Viral Transduction

For lentiviral transduction, open reading frames were cloned into pLenti-CMV-GFP-blast plasmids. Constructs were transfected into 293FT cells together with psPAX2 (Addgene #12260) and pMD2.G (Addgene #12259) using calcium phosphate precipitation. Supernatants containing lentivirus were filtered through a 0.45 μm filter and supplemented with 4 μg/mL polybrene. Successfully transduced cells were selected using 5 μg/mL blasticidin (Thermo Fisher Scientific #R21001).

### Nuclease Assay with Recombinant TREX1

*In vitro* DNA degradation assay was performed as previously described with minor modifications (Zhou et al., 2022). Briefly, 1 μM 100-bp dsDNA (see below for sequence) was incubated with 0.1 μM human TREX1 or TREX1 variants in a 20 μL reaction system (20 mM Tris-HCl pH 7.5, 15 mM NaCl, 135 mM KCl, 5 mM MgCl2, and 1 mg/ml BSA) at 25°C with a time gradient of 5– 30 min. DNA degradation was quenched by adding SDS (final concentration at 0.0167% (w/v)) and EDTA (final concentration at 10 mM) and incubating at 75°C for 15 min. The remaining DNA was separated on a 4% agarose gel using 0.5 × TB buffer (45 mM Tris, 45 mM boric acid) as a running buffer. After DNA electrophoresis, the agarose gel was stained with 0.5x TB buffer (containing 10 μg/mL ethidium bromide) at 25°C for 15 min, followed by de-staining with milli-Q water for an additional 45 min. DNA was visualized by ImageQuant 800 Imaging System and quantified using FIJI (Schindelin et al., 2012).

100-bp dsDNA sense:

5′- ACATCTAGTACATGTCTAGTCAGTATCTAGTGATTATCTAGACATACATCTAGTACATGTCTAGTCAGTATCTAGTGATTATCTAGACATGGACTCATCC -3′

100-bp dsDNA anti-sense:

5′- GGATGAGTCCATGTCTAGATAATCACTAGATACTGACTAGACATGTACTAGATGTATGTCTAGATAATCACTAGATACTGACTAGACATGTACTAGATGT -3′

### Nuclease Assay in Cell Lysates

dsDNA substrate was prepared by annealing oligo 1 (IDT; /5TEX615/GCTAGGCAG) and oligo 2 (IDT; CTGCCTAGC/3IAbRQSp/) in DNA duplex buffer (100 mM KAc, 30 mM HEPES pH 7.5) at a 1:1.15 ratio.

Whole cell lysates were generated by resuspending 3 million cells in 80 μL of assay buffer containing 25 mM HEPES 7.5, 20 mM KCl, 1 mM DTT, 1% Triton X-100, 0.25 mM EDTA, and 10 mM MgCl_2_ supplemented with Complete Mini Protease Inhibitor Cocktail (Invitrogen #11836153001). Cells were lysed by passing the cell resuspension through a 28 G syringe (BD #329461) ten times, incubated on ice for 15 minutes, and then were spun down at 14,000 ✕ *g*, 4°C for 15 minutes to remove pellets. 1:10 dilution of whole cell lysates in assay buffer were used to quantify protein content using Reducing Agent-compatible Pierce BCA Assay Kit (Thermo Fisher Scientific #23250).

2.5 μg (Fig. 1E) or 50 μg (Fig. 3E) of protein was loaded onto a 384-well F-bottom polystyrene microplate (Greiner Bio-One International AG Cat# 784076) with 1 μM dsDNA substrate in assay buffer. The fluorescence intensity (excitation = 570 nm, emission = 615 nm) of the plate was read immediately with Cytation 3 Multi-mode Reader (BioTek) at 25° C for 4 hours every 3 minutes.

### Live-cell Imaging

Cells were plated onto 4-well glass-bottom μ-slide dishes (Ibidi #80427) 24 h before imaging. Five minutes before imaging, media in each well was replaced with FluoroBrite DMEM Imaging Media (Thermo Scientific #A1896701) containing 1 μM ER Tracker Red (Thermo Scientific #E34250) or ER Tracker Green (Thermo Scientific #E34251). Live-cell imaging was performed at room temperature using Nikon Eclipse Ti2-E equipped with CSU-W1 SoRa spinning disk super resolution confocal system, Borealis microadapter, Perfect Focus 4, motorized turret and encoded stage, 5-line laser launch [405 (100 mw), 445 (45 mw), 488 (100 mw), 561 (80 mw), 640 (75 mw)], PRIME 95B Monochrome Digital Camera, and CFI Apo TIRF 60x 1.49 NA objective lens. Images were acquired using NIS-Elements Advanced Research Software on a Dual Xeon Imaging workstation. Adjustment of brightness and contrast were performed using Fiji software. Images were cropped and assembled into figures using Illustrator 2024 (Adobe).

### Immunoblotting

Whole cell lysates were generated by resuspending 1 million cells in RIPA buffer (25 mM Tris-HCl pH 7.6, 150 mM NaCl, 1 % NP-40, 1 % sodium deoxycholate, 0.1 % SDS) supplemented with phosphatase inhibitors (10 mM NaF, 20 mM β-glycerophosphate) and 100 μM phenylmethylsulfonyl fluoride. Cells were lysed by sonication for 15 cycles (high, 30 seconds on, 30 seconds off) using Bioruptor Plus (Diagenode). After a 15-minute incubation on ice and centrifugation (21,000 ✕ *g*, 4 °C for 20 minutes), pellets were removed. 1:10 dilution of whole cell lysates in RIPA were used to quantify protein content using Pierce BCA Assay Kit (Thermo Fisher Scientific #23227). 20 μg protein was loaded per sample into 15-well Novex WedgeWell Tris-Glycine Mini gels (Invitrogen #XP08165BOX). Gels were run at 120 V for 90 minutes and then transferred onto 0.45 μm nitrocellulose membranes (Cytiva #10600002) at 100 V for 60 minutes on ice. Membranes were blocked in Intercept Blocking Buffer (LI-COR #NC1660556). Primary antibodies were diluted (1:4000 for β-actin, 1:1000 for all others) in Intercept T20 (TBS) Antibody Diluent (LI-COR #927-65001) and incubated with membranes overnight at 4 °C on a nutator. Membranes were washed three times in TBST. Secondary antibodies were diluted 1:10,000 in Intercept T20 (TBS) Antibody Diluent and incubated for 1 hour at room temperature on a shaker. After three rounds of washing with TBST and one round of washing with TBS, membranes were scanned using the Odyssey XL infrared imaging scanner (LI-COR).

### 2′3′-cGAMP Quantification

2 million cells were seeded onto 10-cm dishes 24 hours before transfection. Each plate was either transfected with 4 μg herring testes (HT-) DNA or mock-transfected using Lipofectamine 3000 (Thermo Fisher Scientific #L3000075). 24 hours after transfection, cells were harvested, washed with PBS, pelleted, flash-frozen in liquid nitrogen, and stored at –80° C. To quantify 2′3′-cGAMP levels, 2 million cells were resuspended in 200 μL LP2 lysis buffer (20 mM Tris-HCl pH 7.7, 100 mM NaCl, 10 mM NaF, 20 mM β-glycerophosphate, 5 mM MgCl_2_, 0.1 % Triton X-100, 5 % glycerol). Cells were lysed by passing the cell resuspension through a 28 G syringe (BD #329461) ten times, incubated on ice for 15 minutes, and then were spun down at 21,300 *g*, 4° C for 20 minutes to remove pellets. 2′3′-cGAMP levels were quantified using the 2′3′-cGAMP ELISA Kit (Arbor Assays #K067-H5) according to the manufacturer’s instructions. 1:10 dilution of lysates in LP2 buffer were used to quantify protein content using Pierce BCA Assay Kit (Thermo Fisher Scientific #23227). The resulting 2′3′-cGAMP levels were normalized to protein content in each sample.

### RT-qPCR

Total RNA was isolated from 1 million cells using Quick RNA Miniprep Kit (Zymo Research #R1055) according to the manufacturer’s instructions. A DNase I digestion step was included prior to eluting the RNA. cDNA was generated from 1000 ng total RNA using the SuperScript IV First-strand Synthesis System (Invitrogen #18091200) with random hexamer and oligo-(dT) priming. Reverse-transcribed samples were treated with RNase H to remove RNA. qPCR was performed with gene-specific primers (see Key Resources Table) and SYBR Green qPCR Master Mix (Applied Biosystems #A25742). qPCR was performed on QuantStudio 6 (Applied Biosystems), using 10 ng of cDNA and 250 nM of each primer on a MicroAmp 384-well reaction plate (Applied Biosystems #4309849). Relative transcription levels were calculated by normalizing to the geometric mean of *ACTB* and *GAPDH* cycle threshold values.

### Thermal Denaturation Assay

10 µM of purified TREX1 mutant protein and 3✕ SYPRO Orange Protein dye (Life Technologies) were loaded into a 96-well reaction plate, in a 20 µL reaction containing 20 mM Tris-HCl pH 7.5, 75 mM KCl, and 1 mM TCEP. Reactions were incubated with an increasing temperature from 20 to 95° C in a Bio-Rad CFX thermocycler with HEX channel fluorescence measurements taken every 0.5° C, and melting temperature (T_m_) was defined as the temperature at which the half of the maximum fluorescence change occurs.

### Statistical Analysis

Information regarding biological replicates, sample size, and statistical testing is provided in the figure legends.

## ACKNOWLEDGEMENTS

We thank Y. Chen and E. Toufektchan for advice on experimental procedures; C. Krumm for the lysate-based *in vitro* nuclease assay protocol; J. Petrini and J. Tyler for advice; and support from the National Cancer Institute (NCI) (R37CA261183), the Pew Charitable Trusts, the Mary Kay Ash Foundation, and the MSKCC Frank A. Howard Fellowship.

## AUTHOR CONTRIBUTIONS

A.S. and J.M. designed the experiments. A.S. performed most experiments and data analysis. Y.C. identified the AGS-linked P61Q mutation from human genetics data. X.L. and W.Z. performed protein-based *in vitro* nuclease assay and *in vitro* thermal shift assay. A.S. and J.M. wrote the manuscript with input from all authors.

## KEY RESOURCES TABLE

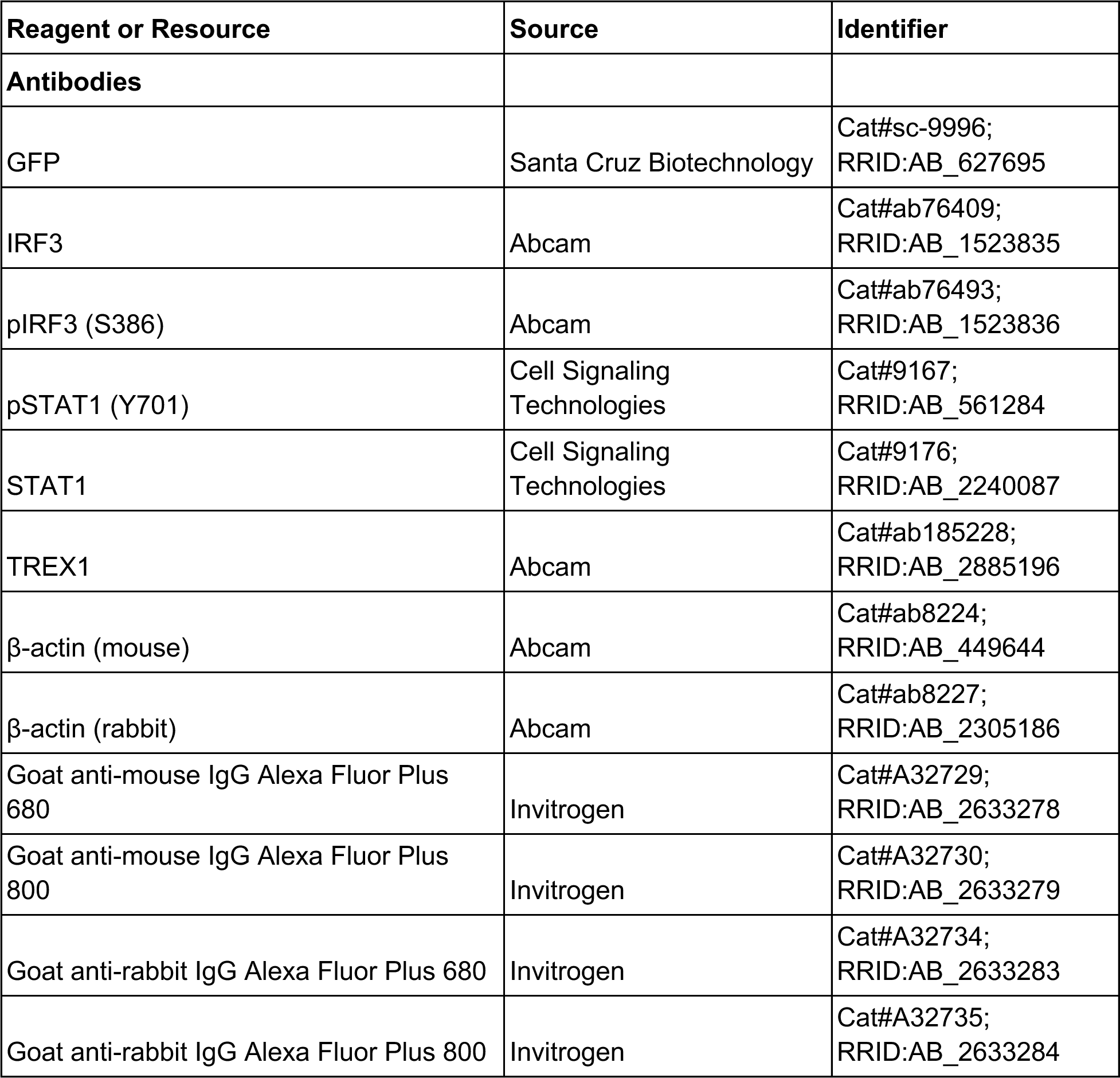
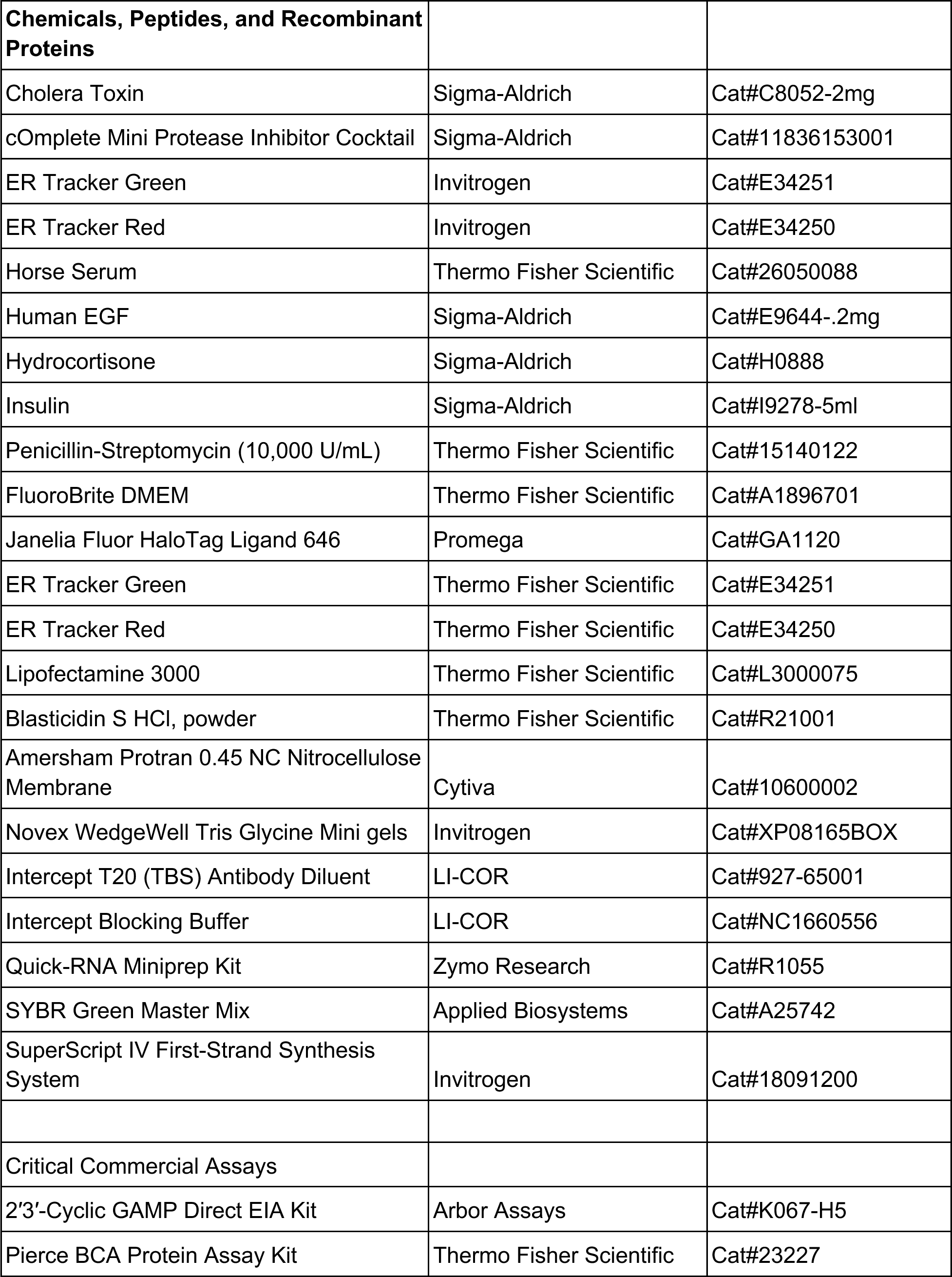
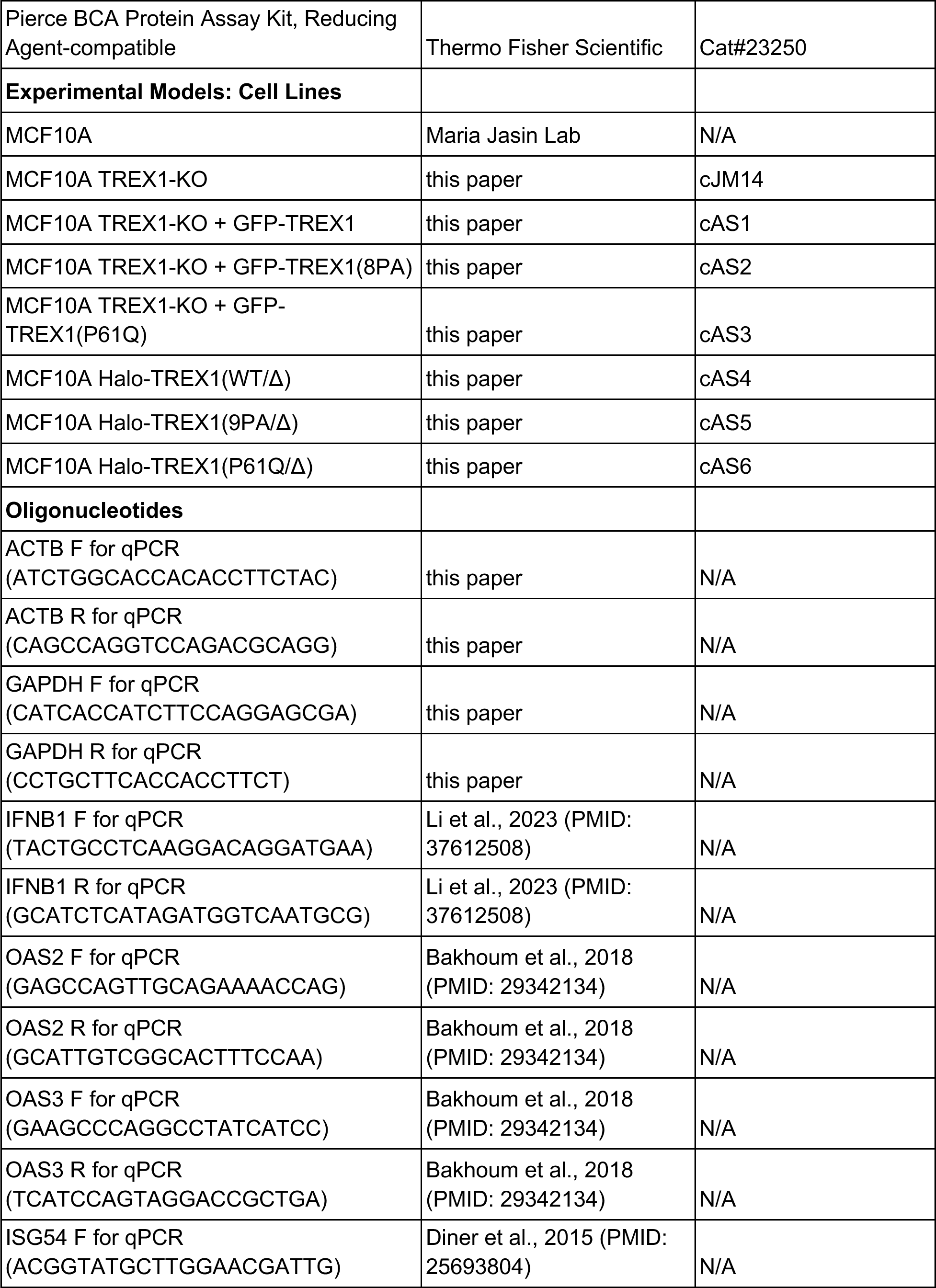
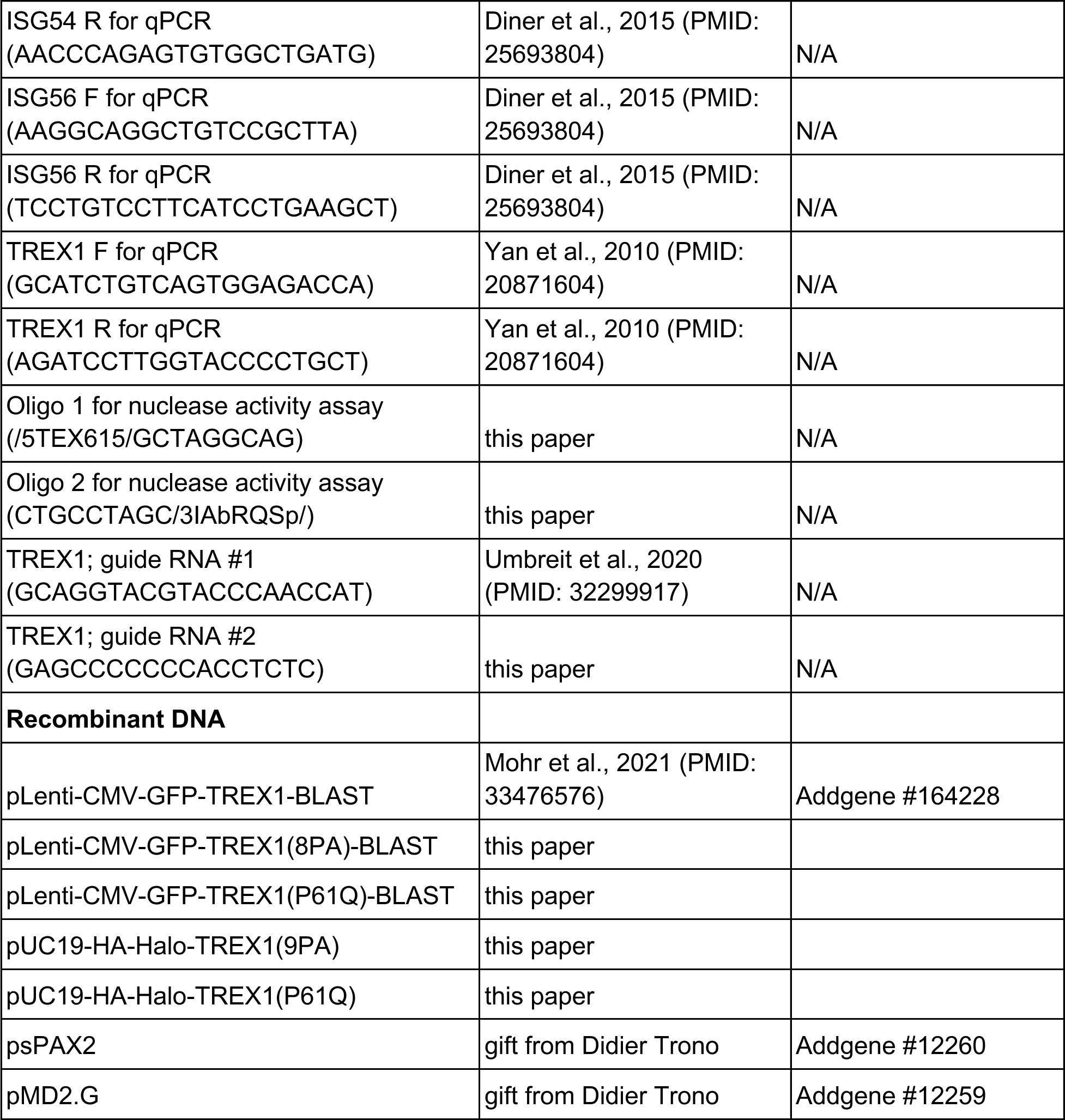

## REFERENCES

Ablasser A, Chen ZJ. 2019. cGAS in action: Expanding roles in immunity and inflammation. Science 363. doi:10.1126/science.aat8657

Ablasser A, Goldeck M, Cavlar T, Deimling T, Witte G, Röhl I, Hopfner K-P, Ludwig J, Hornung V. 2013. cGAS produces a 2’-5’-linked cyclic dinucleotide second messenger that activates STING. Nature 498:380–384.

Ablasser A, Hemmerling I, Schmid-Burgk JL, Behrendt R, Roers A, Hornung V. 2014. TREX1 deficiency triggers cell-autonomous immunity in a cGAS-dependent manner. J Immunol 192:5993–5997.

Adzhubei AA, Sternberg MJE, Makarov AA. 2013. Polyproline-II helix in proteins: structure and function. J Mol Biol 425:2100–2132.

Ahn J, Ruiz P, Barber GN. 2014. Intrinsic self-DNA triggers inflammatory disease dependent on STING. J Immunol 193:4634–4642.

Broadbent DG, Barnaba C, Perez GI, Schmidt JC. 2023. Quantitative analysis of autophagy reveals the role of ATG9 and ATG2 in autophagosome formation. J Cell Biol 222. doi:10.1083/jcb.202210078

Brucet M, Querol-Audí J, Serra M, Ramirez-Espain X, Bertlik K, Ruiz L, Lloberas J, Macias MJ, Fita I, Celada A. 2007. Structure of the dimeric exonuclease TREX1 in complex with DNA displays a proline-rich binding site for WW Domains. J Biol Chem 282:14547–14557.

Chowdhury D, Beresford PJ, Zhu P, Zhang D, Sung J-S, Demple B, Perrino FW, Lieberman J. 2006. The exonuclease TREX1 is in the SET complex and acts in concert with NM23-H1 to degrade DNA during granzyme A-mediated cell death. Mol Cell 23:133–142.

Crow YJ, Chase DS, Lowenstein Schmidt J, Szynkiewicz M, Forte GMA, Gornall HL, Oojageer A, Anderson B, Pizzino A, Helman G, Abdel-Hamid MS, Abdel-Salam GM, Ackroyd S, Aeby A, Agosta G, Albin C, Allon-Shalev S, Arellano M, Ariaudo G, Aswani V, Babul-Hirji R, Baildam EM, Bahi-Buisson N, Bailey KM, Barnerias C, Barth M, Battini R, Beresford MW, Bernard G, Bianchi M, Billette de Villemeur T, Blair EM, Bloom M, Burlina AB, Carpanelli ML, Carvalho DR, Castro-Gago M, Cavallini A, Cereda C, Chandler KE, Chitayat DA, Collins AE, Sierra Corcoles C, Cordeiro NJV, Crichiutti G, Dabydeen L, Dale RC, D’Arrigo S, De Goede CGEL, De Laet C, De Waele LMH, Denzler I, Desguerre I, Devriendt K, Di Rocco M, Fahey MC, Fazzi E, Ferrie CD, Figueiredo A, Gener B, Goizet C, Gowrinathan NR, Gowrishankar K, Hanrahan D, Isidor B, Kara B, Khan N, King MD, Kirk EP, Kumar R, Lagae L, Landrieu P, Lauffer H, Laugel V, La Piana R, Lim MJ, Lin J-PS-M, Linnankivi T, Mackay MT, Marom DR, Marques Lourenço C, McKee SA, Moroni I, Morton JEV, Moutard M-L, Murray K, Nabbout R, Nampoothiri S, Nunez-Enamorado N, Oades PJ, Olivieri I, Ostergaard JR, Pérez-Dueñas B, Prendiville JS, Ramesh V, Rasmussen M, Régal L, Ricci F, Rio M, Rodriguez D, Roubertie A, Salvatici E, Segers KA, Sinha GP, Soler D, Spiegel R, Stödberg TI, Straussberg R, Swoboda KJ, Suri M, Tacke U, Tan TY, te Water Naude J, Wee Teik K, Thomas MM, Till M, Tonduti D, Valente EM, Van Coster RN, van der Knaap MS, Vassallo G, Vijzelaar R, Vogt J, Wallace GB, Wassmer E, Webb HJ, Whitehouse WP, Whitney RN, Zaki MS, Zuberi SM, Livingston JH, Rozenberg F, Lebon P, Vanderver A, Orcesi S, Rice GI. 2015. Characterization of human disease phenotypes associated with mutations in TREX1, RNASEH2A, RNASEH2B, RNASEH2C, SAMHD1, ADAR, and IFIH1. Am J Med Genet A 167A:296–312.

Crow YJ, Manel N. 2015. Aicardi-Goutières syndrome and the type I interferonopathies. Nat Rev Immunol 15:429–440.

Crow YJ, Stetson DB. 2022. The type I interferonopathies: 10 years on. Nat Rev Immunol 22:471–483.

De Silva U, Choudhury S, Bailey SL, Harvey S, Perrino FW, Hollis T. 2007. The crystal structure of TREX1 explains the 3′ nucleotide specificity and reveals a polyproline II helix for protein partnering. J Biol Chem 282:10537–10543.

Diner EJ, Burdette DL, Wilson SC, Monroe KM, Kellenberger CA, Hyodo M, Hayakawa Y, Hammond MC, Vance RE. 2013. The innate immune DNA sensor cGAS produces a noncanonical cyclic dinucleotide that activates human STING. Cell Rep 3:1355–1361.

Galluzzi L, Vanpouille-Box C, Bakhoum SF, Demaria S. 2018. SnapShot: CGAS-STING Signaling. Cell 173:276–276.e1.

Gao D, Li T, Li X-D, Chen X, Li Q-Z, Wight-Carter M, Chen ZJ. 2015. Activation of cyclic GMP-AMP synthase by self-DNA causes autoimmune diseases. Proc Natl Acad Sci U S A 112:E5699–705.

Gao P, Ascano M, Zillinger T, Wang W, Dai P, Serganov AA, Gaffney BL, Shuman S, Jones RA, Deng L, Others. 2013. Structure-function analysis of STING activation by c [G (2′, 5′) pA (3′, 5′) p] and targeting by antiviral DMXAA. Cell 154:748–762.

Gray EE, Treuting PM, Woodward JJ, Stetson DB. 2015. Cutting Edge: cGAS Is Required for Lethal Autoimmune Disease in the Trex1-Deficient Mouse Model of Aicardi–Goutières Syndrome. The Journal of Immunology 195:1939–1943.

Grieves JL, Fye JM, Harvey S, Grayson JM, Hollis T, Perrino FW. 2015. Exonuclease TREX1 degrades double-stranded DNA to prevent spontaneous lupus-like inflammatory disease. Proc Natl Acad Sci U S A 112:5117–5122.

Lee-Kirsch MA, Gong M, Chowdhury D, Senenko L, Engel K, Lee Y-A, de Silva U, Bailey SL, Witte T, Vyse TJ, Kere J, Pfeiffer C, Harvey S, Wong A, Koskenmies S, Hummel O, Rohde K, Schmidt RE, Dominiczak AF, Gahr M, Hollis T, Perrino FW, Lieberman J, Hübner N. 2007. Mutations in the gene encoding the 3’-5’ DNA exonuclease TREX1 are associated with systemic lupus erythematosus. Nat Genet 39:1065–1067.

Lehtinen DA, Harvey S, Mulcahy MJ, Hollis T, Perrino FW. 2008. The TREX1 double-stranded DNA degradation activity is defective in dominant mutations associated with autoimmune disease. J Biol Chem 283:31649–31656.

Liu S, Cai X, Wu J, Cong Q, Chen X, Li T, Du F, Ren J, Wu Y-T, Grishin NV, Chen ZJ. 2015. Phosphorylation of innate immune adaptor proteins MAVS, STING, and TRIF induces IRF3 activation. Science 347:aaa2630.

Mazur DJ, Perrino FW. 2001. Excision of 3′ Termini by the Trex1 and TREX2 3′→ 5′ Exonucleases characterization of the recombinant proteins. J Biol Chem 276:17022–17029.

Mohr L, Toufektchan E, von Morgen P, Chu K, Kapoor A, Maciejowski J. 2021. ER-directed TREX1 limits cGAS activation at micronuclei. Mol Cell. doi:10.1016/j.molcel.2020.12.037

Rice GI, Melki I, Frémond M-L, Briggs TA, Rodero MP, Kitabayashi N, Oojageer A, Bader-Meunier B, Belot A, Bodemer C, Quartier P, Crow YJ. 2017. Assessment of Type I Interferon Signaling in Pediatric Inflammatory Disease. J Clin Immunol 37:123–132.

Rice GI, Rodero MP, Crow YJ. 2015. Human disease phenotypes associated with mutations in TREX1. J Clin Immunol 35:235–243.

Rice G, Newman WG, Dean J, Patrick T, Parmar R, Flintoff K, Robins P, Harvey S, Hollis T, O’Hara A, Herrick AL, Bowden AP, Perrino FW, Lindahl T, Barnes DE, Crow YJ. 2007a. Heterozygous mutations in TREX1 cause familial chilblain lupus and dominant Aicardi-Goutieres syndrome. Am J Hum Genet 80:811–815.

Rice G, Patrick T, Parmar R, Taylor CF, Aeby A, Aicardi J, Artuch R, Montalto SA, Bacino CA, Barroso B, Baxter P, Benko WS, Bergmann C, Bertini E, Biancheri R, Blair EM, Blau N, Bonthron DT, Briggs T, Brueton LA, Brunner HG, Burke CJ, Carr IM, Carvalho DR, Chandler KE, Christen H-J, Corry PC, Cowan FM, Cox H, D’Arrigo S, Dean J, De Laet C, De Praeter C, Dery C, Ferrie CD, Flintoff K, Frints SGM, Garcia-Cazorla A, Gener B, Goizet C, Goutieres F, Green AJ, Guet A, Hamel BCJ, Hayward BE, Heiberg A, Hennekam RC, Husson M, Jackson AP, Jayatunga R, Jiang Y-H, Kant SG, Kao A, King MD, Kingston HM, Klepper J, van der Knaap MS, Kornberg AJ, Kotzot D, Kratzer W, Lacombe D, Lagae L, Landrieu PG, Lanzi G, Leitch A, Lim MJ, Livingston JH, Lourenco CM, Lyall EGH, Lynch SA, Lyons MJ, Marom D, McClure JP, McWilliam R, Melancon SB, Mewasingh LD, Moutard M-L, Nischal KK, Ostergaard JR, Prendiville J, Rasmussen M, Rogers RC, Roland D, Rosser EM, Rostasy K, Roubertie A, Sanchis A, Schiffmann R, Scholl-Burgi S, Seal S, Shalev SA, Corcoles CS, Sinha GP, Soler D, Spiegel R, Stephenson JBP, Tacke U, Tan TY, Till M, Tolmie JL, Tomlin P, Vagnarelli F, Valente EM, Van Coster RNA, Van der Aa N, Vanderver A, Vles JSH, Voit T, Wassmer E, Weschke B, Whiteford ML, Willemsen MAA, Zankl A, Zuberi SM, Orcesi S, Fazzi E, Lebon P, Crow YJ. 2007b. Clinical and molecular phenotype of Aicardi-Goutieres syndrome. Am J Hum Genet 81:713–725.

Rodero MP, Decalf J, Bondet V, Hunt D, Rice GI, Werneke S, McGlasson SL, Alyanakian M-A, Bader-Meunier B, Barnerias C, Bellon N, Belot A, Bodemer C, Briggs TA, Desguerre I, Frémond M-L, Hully M, van den Maagdenberg AMJM, Melki I, Meyts I, Musset L, Pelzer N, Quartier P, Terwindt GM, Wardlaw J, Wiseman S, Rieux-Laucat F, Rose Y, Neven B, Hertel C, Hayday A, Albert ML, Rozenberg F, Crow YJ, Duffy D. 2017. Detection of interferon alpha protein reveals differential levels and cellular sources in disease. J Exp Med 214:1547–1555.

Schindelin J, Arganda-Carreras I, Frise E, Kaynig V, Longair M, Pietzsch T, Preibisch S, Rueden C, Saalfeld S, Schmid B, Tinevez J-Y, White DJ, Hartenstein V, Eliceiri K, Tomancak P, Cardona A. 2012. Fiji: an open-source platform for biological-image analysis. Nat Methods 9:676–682.

Stetson DB, Ko JS, Heidmann T, Medzhitov R. 2008. Trex1 prevents cell-intrinsic initiation of autoimmunity. Cell 134:587–598.

Sudol M, Chen HI, Bougeret C, Einbond A, Bork P. 1995. Characterization of a novel protein-binding module--the WW domain. FEBS Lett 369:67–71.

Wolf C, Rapp A, Berndt N, Staroske W, Schuster M, Dobrick-Mattheuer M, Kretschmer S, König N, Kurth T, Wieczorek D, Kast K, Cardoso MC, Günther C, Lee-Kirsch MA. 2016. RPA and Rad51 constitute a cell intrinsic mechanism to protect the cytosol from self DNA. Nat Commun 7:11752.

Yan N. 2017. Immune Diseases Associated with TREX1 and STING Dysfunction. J Interferon Cytokine Res 37:198–206.

Zhou W, Mohr L, Maciejowski J, Kranzusch PJ. 2021. cGAS phase separation inhibits TREX1-mediated DNA degradation and enhances cytosolic DNA sensing. Mol Cell 81:739–755.e7.

Zhou W, Richmond-Buccola D, Wang Q, Kranzusch PJ. 2022. Structural basis of human TREX1 DNA degradation and autoimmune disease. Nat Commun 13:4277.

